# Deviation from mendelian transmission of autosomal SNPs can be used to estimate germline mutations in humans exposed to ionizing radiation

**DOI:** 10.1101/2020.05.18.101816

**Authors:** Hugo Pereira Leite Filho, Irene Plaza Pinto, Lorraynne Guimarães Oliveira, Emília Oliveira Alves Costa, Alex Silva da Cruz, Daniela de Melo e Silva, Claudio Carlos da Silva, Alexandre Rodrigues Caetano, Aparecido Divino da Cruz

## Abstract

We aimed to estimate the rate of germline mutations in the offspring of individuals accidentally exposed to Cesium-137 ionizing radiation. Performed analysis considered two distinct groups: a group males and females accidentally exposed to low doses of ionizing radiation from Cs^137^, the case group, and a control group of non-exposed participants. The case group included 37 participants (11 couples and 15 children born after the accident). The dose absorbed by exposed participants ranged from 0.2 to 0.5 Gray. The control group included 15 families from the state of Goiás, with no history of radiation exposure. DNA samples from peripheral blood were analyzed with the Affymetrix GeneChip^®^ CytoScanHD^™^ to assess *de novo* SNP-type mutations. A set of scripts previously developed was used to detect *de novo* mutations by comparing parent and offspring genotypes in each SNP marker. Overall numbers of observed Mendelian deviations were statistically significant between the exposed and control groups. Offspring from the population accidentally exposed to low IR doses showed ∼ 46.5% more *de novo* Mendelian deviations than the control group. Parent-of-origin and type of nucleotide substitution were also inferred. Estimates of age-adjusted *de novo* germline mutation rates were obtained and correlated to Cs-137 radiation dose exposure to evaluate the usefulness of the rate of Mendelian deviations observed in polymorphic SNPs as a biomarker for exposure. This proved useful in a retrospective estimation of the rate of *de novo* germline mutations in a human population accidentally exposed to low doses of radiation from Cs-137. Obtained results suggest that observed burden of germline mutations identified in offspring could potentially be a useful biomarker to estimate levels of parental exposure to ionizing radiation.

## Introduction

In 1987, a series of unexpected events resulted in a major radiological accident in Goiânia, capital of the State of Goiás (Brazil), causing human, animal, plant and environmental exposure to ionizing radiation (IR) and contamination by Cesium-137 [1]. In the aftermath, 249 people were exposed to IR from Cs-137. For some people, individual exposure resulted from internal and external contamination by the radioactive salt, while others were exposed to radiation emitted by the decay of Cs-137. In some cases, people were both exposed to radiation and contaminated by the radionuclide. During the accident, individual absorbed doses of radiation ranged from 0 to 7 Gy, resulting in four fatalities during the acute phase of the accident [2,3].

Following the accident, the Goiânia exposed population has been extensively monitored using genetic biomarkers, as they have been shown to be efficient biomarkers of exposure to gamma rays [4]. However, each biomarker tends to reveal a distinct biological phenomenon in the exposed cells, mostly associated with DNA repair and how cells physiologically coped to survive a specific insult. In this context, our group and others have identified somatic mutations on DNA sequence data from glycophorin A [5] and HPRT [6], general [7,8] and specific chromosomal aberrations, identified by FISH [9] and BCL2/J(H) translocation assays [10], micronucleus presence [11], and assessment of autoimmune markers data (da Cruz et al. *unpublished data*) in the cohort accidently exposed to Cs-137 IR. Moreover, in order to understand the effect of IR on the induction of germ line mutation rates on STR markers [12] have also been determined. More recently, CNVs have been used as biomarkers for parental exposure to demonstrate the effect of low absorbed doses of IR on germline mutations in the cohort’s offspring conceived after the accident [13].

Absorbed dose of IR relate to the estimated quantity of energy deposited in mater per unit of mass. Thus, it can be used as an indirect measurement of the harmful biological effect of the radioactive energy on the cellular system. It is calculated by estimating the concentration of energy from radiation exposure deposited in each organ, using a reference value, the type of radiation and the potential for radiation-related mutagenic changes in each organ or tissue [14].

The exposure of cells to IR delays the normal progression of the cell cycle [15-17], initially observed as a passive cellular response resulting from of the induction of DNA damage in the exposed cells. The irradiated cell must adapt to the insult and facilitate DNA repair processes, especially fixing double-strand breaks, the most common damage after DNA exposure to IR [18-22].

The mutagenic effects of IR on the human germ line cells are of concern, as they lead to the accumulation of mutations in the offspring of irradiated parents, which amounts to an increase in the mutational burden [23]. Despite numerous studies, little is known about the genetic effects of low doses of radiation from low linear energy transfer gamma radiation exposure in humans. Most of the consolidated evidence comes from extrapolation of the induction of germline mutations in mammals, often rat and mouse models [24,25].

Advances in the techniques of genomic analysis have greatly increased the volume of nucleotide sequence data, enabling the identification of thousands of SNPs (single-nucleotide polymorphisms). Variations in SNPs are important to determine genotypic and phenotypic relationships, within and between species and populations, and also to identify variants related to genetic diseases in humans and animals [26]. In this context, genomic analysis can be a useful tool to study and understand the effects of IR exposure on animals and humans [13,27].

In recent decades, several genotyping technologies have been developed to characterize SNPs all producing genotype matrices with hundreds of thousands of datapoints. Algorithms based on parametric and nonparametric statistical models have been used to determine the genotype of each SNP from the fluorescence signal intensity of marked probes, which are scanned, captured, and arranged in a matrix format [28,29]. One of the outstanding commercially available SNP arrays is the GeneChip^®^ CytoScanHD^™^ (Thermo Fisher, Massachusetts, USA), considered to be a high-density matrix, with about 750,000 SNP markers with average genotyping accuracy >99% [30].

In the aforementioned context, the general objective of the current study was to identify Mendelian deviations (MD) in genome-wide autosomal SNP data from a cohort of people conceived after parental exposure to Cs-137 IR, and a group of non-exposed people from the same geographical area, and to evaluate if the observed burden of germline mutations identified in offspring could be a potentially useful biomarker to estimate levels of parental exposure to ionizing radiation.

## Material and Methods

### Sample collection, processing, and genotyping

The experiment was designed as a case-control observational study. The group of case subjects consisted of 11 families (Table 1), of whom at least one parent was accidentally exposed to IR during the Cs-137 accident, totaling 37 participants (11 fathers, 11 mothers, and 15 children conceived after the accident). The absorbed doses for the exposed parents ranged from 0.2 to 0.5 Gy. A group of participants with no prior history of IR exposure was used as a control. Control samples were obtained from 15 families living in Goiânia, comprised of 15 fathers, 15 mothers, and 15 children also conceived after 1987. A total of 82 subjects were used in the study. DNA samples from all subjects were analyzed with the GeneChip^®^ CytoScanHD^™^ chip (Thermo Fisher, Massachusetts, USA). Cases and controls participated voluntarily in the study, which was approved by the ethics committee on research with humans from the Pontifical Catholic University of Goiás (PUC-Goiás) – CAAE number 49338615.2.0000.0037. At the time of blood collection, the participants answered a questionnaire and signed an informed consent and/or authorization form, where pertinent. A total of 10 mL of peripheral blood in EDTA was voluntarily donated by all participants. Total genomic DNA was isolated from whole blood using Illustra blood genomicPrep Mini Spin Kit® (GE Healthcare) and stored at −20°C. The remaining biological material was stored for future studies, according to CNS Resolution 441/11.

**Table 1.** Variables generated by ChAS® and considered by SIPO to identify MDs.

| **Marker** | **SNP marker identifier** |
| --- | --- |
| **Genotyping** | Genotyping call. Biallelic information. Three possible combinations: AA/BB/AB |
| **Trust Value** | Confidence value for each genotyping call |
| **Sign A** | Gross sign value for sign A on the marker |
| **Sign B** | Gross sign value for sign B on the marker |
| **Nitrogen Base** | Call from the nitrogen base. Biallelic information. Ten possible combinations: AA\|AC\|AG\|AT\|CC\|CG\|CT\|GG\|TT |
| **dbSNP** | SNP identifier record in the NCBI dbSNP database |
| **Chromosome** | Autosome associated with the marker |
| **Chromosome Position** | SNP locus on the chromosome |

SNP genotypes were generated with ChAS^©^ with filter parameters set at 15 and 8 markers to detect genomic gains and losses, respectively, distributed with a mean of ≤ 2,000 pb, limited to fragments ≥1pb. The hg19 version of the human genome, contained in the Genome Browser of the University of California, Santa Clara, CA, USA, was also used for genotype analyses.

SNP data quality control was performed according to guidelines from the manufacturer. Genotypes were filtered based on call confidence levels - all genotypes with confidence levels below 5×10^−2^ were excluded from further analysis. In a final step, all markers showing “no call” or “null” in one or more samples were removed from the dataset. Therefore, only markers with quality-controlled genotypes in all samples were considered in the remaining analysis.

### Principle Component Analysis

Principle Component Analysis (PCA) methods were used to assess whether participants in the case and control groups came from the same genetic population. This step was also included to assess whether individual sample quality effects may have generated spurious results. SNPs with minor allele frequencies (MAF) below 0.01 were removed from the dataset. The final dataset contained about 522K SNPs. Data pruning was performed using the PLINK package [31] to generate a subset of markers for PCA analysis using the following parameters: window size of 500 SNP with a step size of 5 SNP, using an r^2^ threshold of 0.1. The resulting subset of SNPs was used to estimate principal components for each test group using PLINK. PCA plots were generated with PLINK.

### Analysis of genotyping data to identify Mendelian Deviations

MDs were identified with a set of previously developed Perl scripts and R libraries [32] termed SIPO (Scripts for Inference of Parental Origin, Leite Filho et al. *submitted*), to mine SNP data in MySql^©^ format. The SIPO pipeline is listed in Fig 1. Parent genotypes were compared with respective offspring marker genotypes. Sex chromosome data were excluded from the analysis, as X-linked data showed elevated noise and Y-specific regions had low marker coverage. Table 1 shows all data variables considered by SIPO.

**Fig 1.**
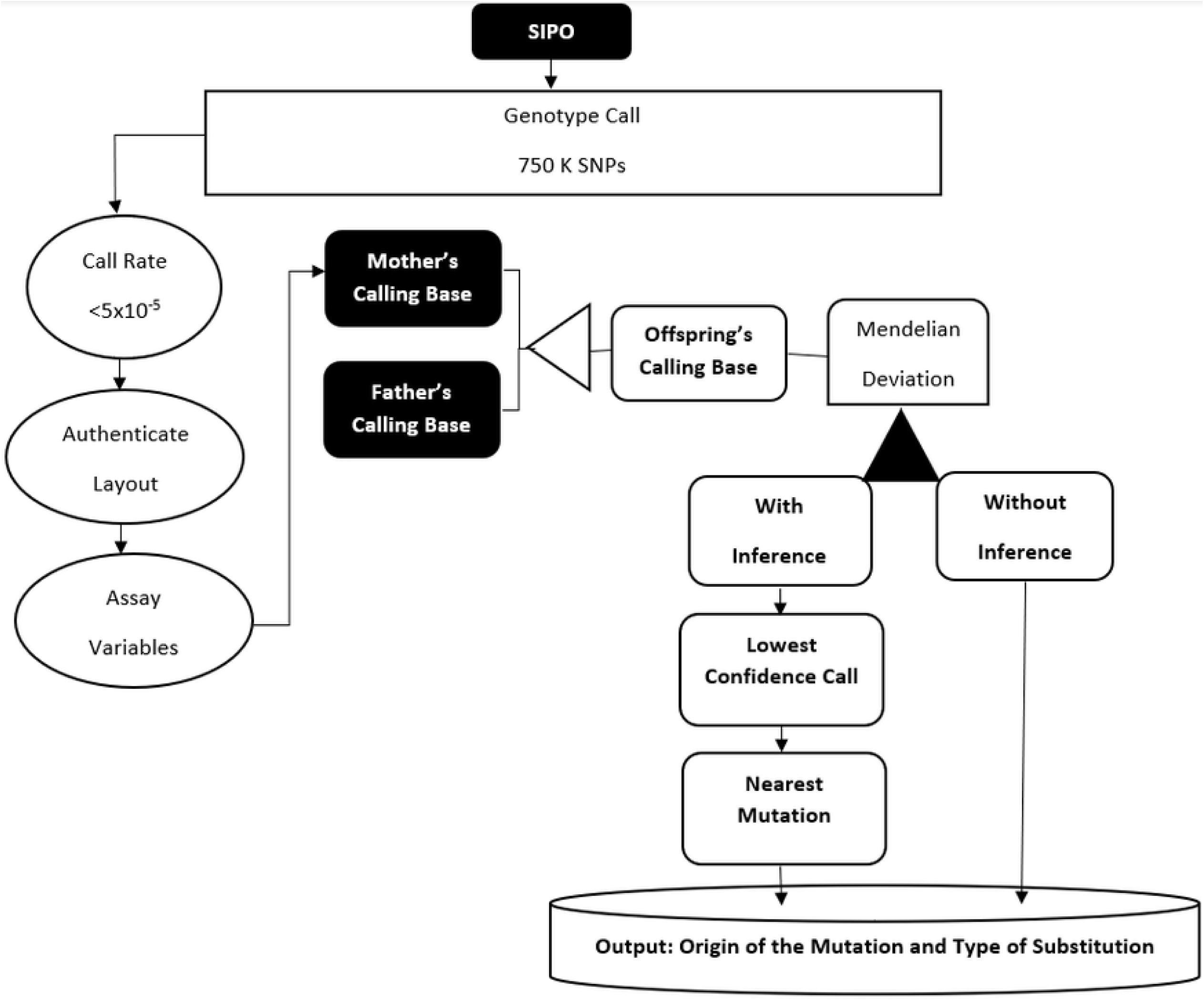
Steps performed by SIPO to infer de novo mutations. Deduct the parental origin of the MD, indicate the type of substitution and generate the estimated rate of Mendelian deviation in the offspring of people accidentally exposed to Cs-137 ionizing radiation.

First, SIPO validated the .CYCHP file generated by ChAS^®^, then SIPO identified trio variables and started to generate inferences for *de novo* mutations, using *a posteriori* markers. Executed steps allowed to determine nucleotide substitution type in addition to inferring the parent of origin of the MD observed in the offspring. Derived information was loaded into a MySQL database and R scripts were used to perform linear regression, clustering and PCA with the resulting data (Fig 2).

**Fig 2.**
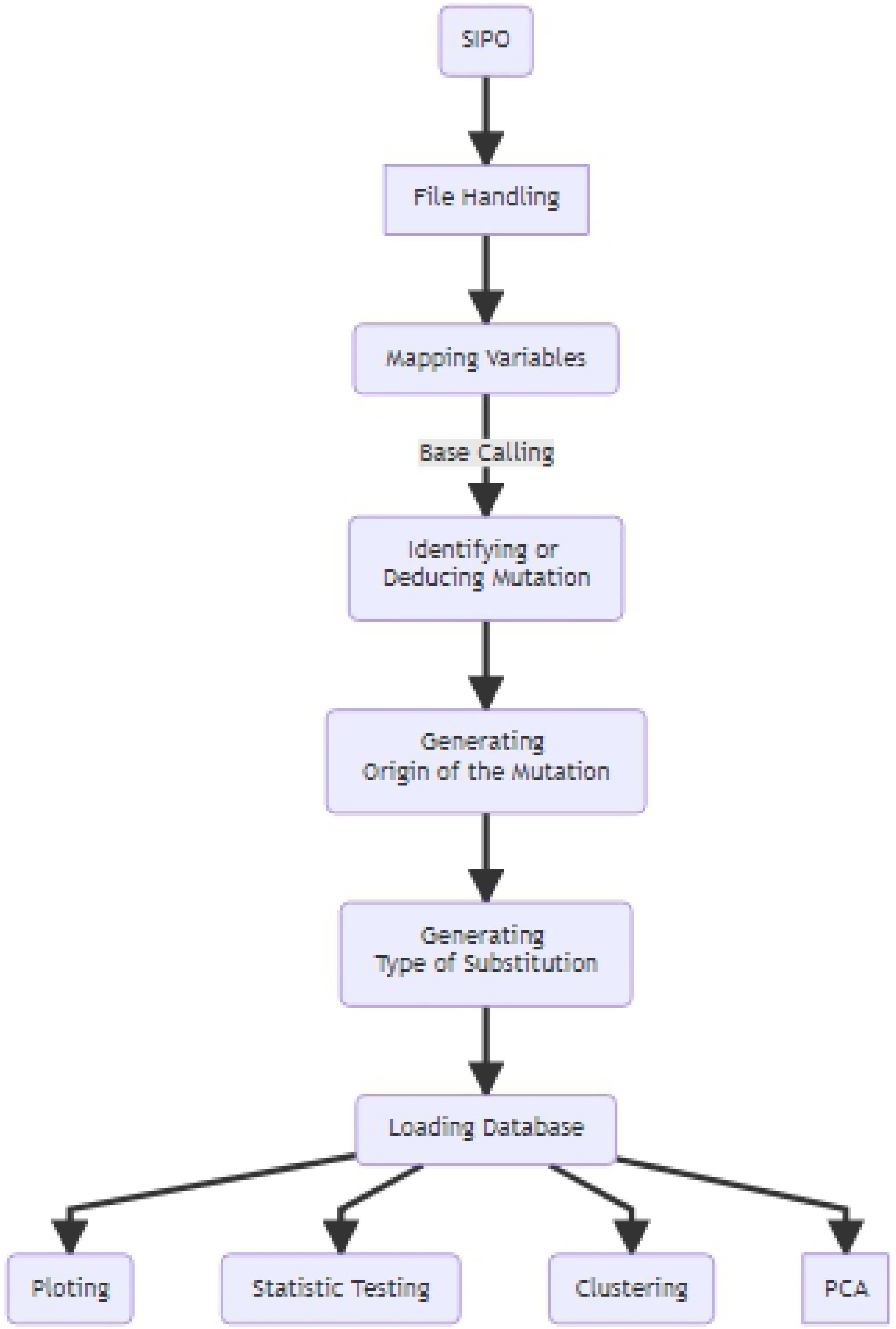
Workflow of the SIPO. Steps performed to infer de novo mutations from SNPs obtained from a genotyping array.

In some situations, it was not possible to identify the parental origin of a SNP by traditional methods. To solve this challenge, two deductions were incorporated into SIPO. The first deduction came from the lower trust value of ChAS^®^ for parents. The second consisted of inferring the MD according to the parental origin of the nearest variant SNP. The Euclidean distance technique was used to calculate differences between positions. Thus, the origin of the mutation was attributed to the parent whose SNP had the lowest confidence value and to the parent who transmitted that chromosomal segment to the child.

MD estimates were used to estimate the germline mutation rate (TM_DM_) in the offspring, using equation 1: [13,33]

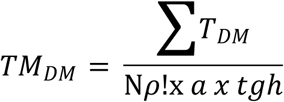

Where ΣT_DM_ = Total MD; Nρ! Is the number of paternal meiosis; a is a biallelic locus (2); tgh is the haploid genome size (2.9 × 10^9^) according to the assembly of the human reference sequence (GRCh37/hg19).

Equation 2 was used to estimate the number of paternal meiosis for cases and controls by age of the father at the time of conception of offspring:

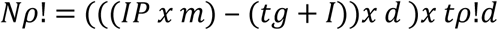

Where Nρ! Is the estimated number of paternal meiosis; IP is the Father’s age at conception (years); m is months/year; tg is gestation time (months); I is mean age of sperm production onset in adolescence, estimated at 12 years old in humans (144 months); d is total number of days in the month; tρ!d is total mean number of estimated meiosis per day in males (25 x 10^6^).

In the present study, all statistical tests were performed considering a 95 % confidence interval and 5 % significance level. The statistical tests used were the Shapiro-Wilk test, regression analysis, Cohen-d [34-37], Tukey, clustering [38], and principal component analysis [39,40]. The R statistical package [32] was used in all analyses.

## Results and discussion

This study used SNP genotypes from a cohort of offspring born to parents accidentally exposed to Cs-137 to estimate germline mutations in humans exposed to low doses of ionizing radiation. Transgenerational and retrospective DNA analysis was used to estimate the contribution of the exposure to the burden of SNP mutations in the offspring. We estimated a 46.5 % increase in the TM_DM_ of the offspring when compared to controls. General data regarding the parents and offspring included in the study are provided in Table 2. The current study establishes a pioneering application of SNP data analysis to identify MD and estimate germline mutations in the offspring of humans accidentally exposed to low absorbed doses of IR. Current findings corroborate our first study reporting the usefulness of small CNVs to estimate de novo human germline mutation rates in the same subjects [13].

**Table 2.** Descriptive data of the case and control groups for parental and F1 generations in the study of the effect of IR exposure on the induction of germline mutations in humans.

| Generation | Variables |  | Cases | Control |
| --- | --- | --- | --- | --- |
| Parental | N |  | 15 | 15 |
|  | Age range (years) |  | 16 – 56 | 19 – 55 |
| | Mean age at conception (years ± SD) | Paternal | 31.4 (12.3) | 32.2 (12.8) |
|  |  | Maternal | 26.4 (5.1) | 27.5 (9.8) |
|  | Absorbed dose (Gy) |  | 0.2 – 0.5 | 0 |
| F1 | N |  | 15 | 15 |
|  | Age range (years) |  | 2 – 20 | 0.85 – 26 |
| | Mean age (years ± SD) | | 14.1 (6.1) | 10.5 (8.5) |
|  | Sex ratio (Male:Female) |  | 8:7 | 10:5 |
| | Mean $TM_{DM}$ (± SD) | | $6.3 \times 10^{-4}$<br>($\pm 2.0 \times 10^{-4}$) | $4.3 \times 10^{-4}$<br>($\pm 9.2 \times 10^{-5}$) |

Data from the study on the induction of Mendelian errors in offspring autosomes of people accidentally exposed to low doses of Cs-137 ionizing radiation. SD = Standard Deviation.

PCA results using a subset of LD-pruned data (522 Kb SNPs) indicates subjects included in case and control groups belong to the same population and that there are no additional confounding factors associated with the test groups other than exposure to Cs-137 (Fig 3). Therefore, the TM_DM_ could be compared between groups, even with a reduced sample size in each group. Observed MDs followed a normal distribution (*p = 0*.5592) and were all included in subsequent statistical analyses. The lowest individual numbers of MDs were 972 and 682, and the highest were 2,875 and 1,635 for the case and control groups, respectively (data not shown). The mean TM_DM_ was approximately 46.5% higher for cases than for controls. SNPs with observed MDs are randomly distributed on the genome. Most MDs were observed in a single trio (60%), while 27%, 9% and 4% MDs were observed in two, three and four trios, respectively (range: 1 – 15).

**Fig 3.**
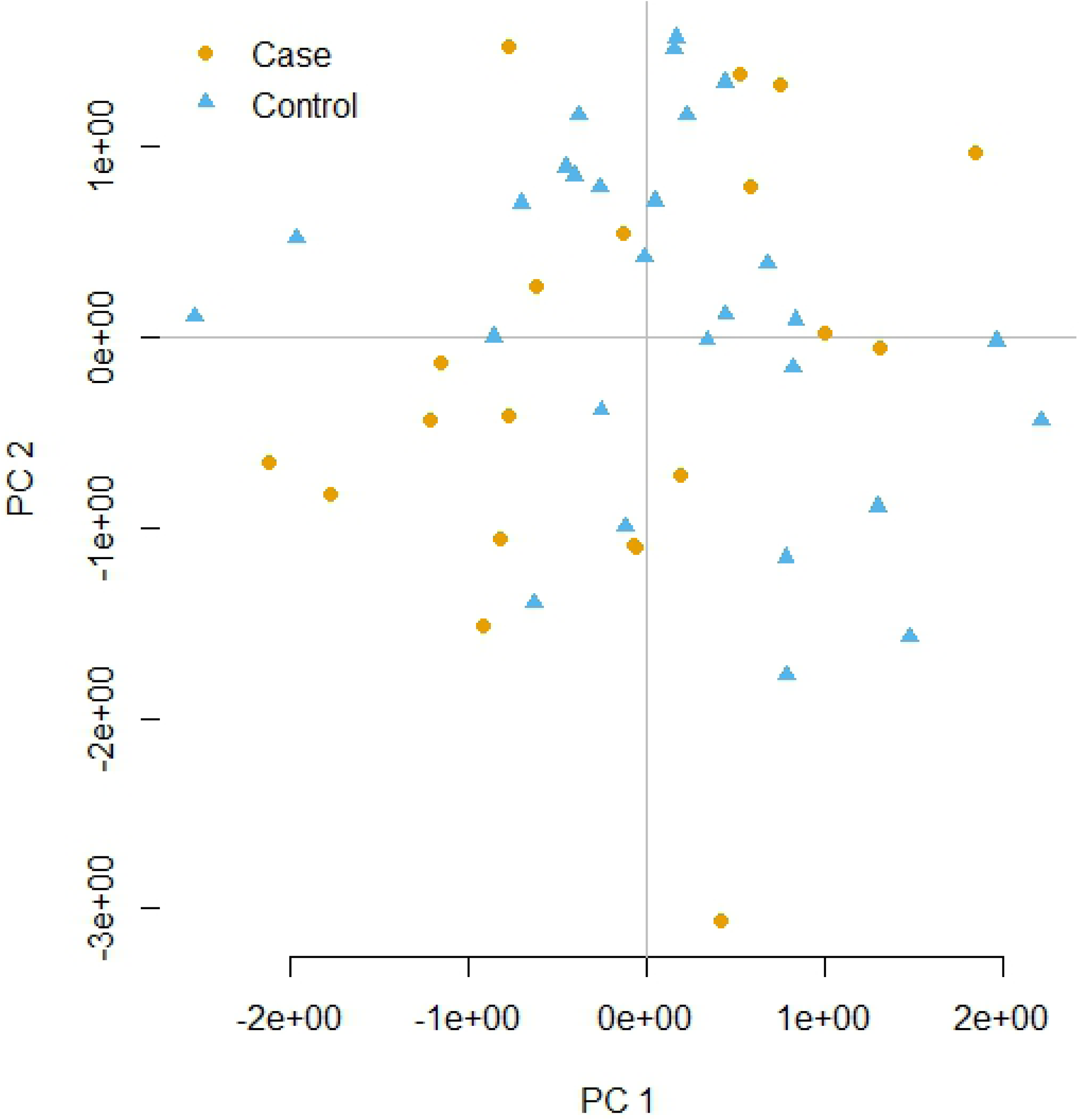
PCA with 2.7K SNP. The variables contained in the PCA represent the standardized relationship matrix by variance, multidimensional sizing (MDS) based on Hamming distances.

Considering the number of observed MDs, adjusted according to parental age, it was possible to estimate the mean TM_DM_ as 6.3 x 10^−4^ and 4.3 x 10^−4^ for case and control groups, respectively. In this study, mutation burden was defined by the number of *de novo* base substitutions in an assayed SNP of a child born to a parent exposed to ionizing radiation. In this context, the germline mutation burden represented by MDs (TM_DMβ_) was significantly higher (p < 2 x 10^−3^) in the offspring of exposed individuals than in the Goiânia control population. In this context, TM_DM_ was a quantifiable and useful variable to estimate the parental contribution to the mutational burden of their children, as a consequence of transmitting non-deleterious mutations. These errors could have been corrected by the DNA repair systems, fixed in the parental germ lines and transmitted to the offspring (Fig 4). The F test, to evaluate MD frequencies in the test groups, showed that the number of observed MDs were significantly different (F *=* 4.80; p < 5 x 10^−3^). Mean MD numbers in the offspring of the case and control groups are shown in Fig 4A. Representation of the number of MD observed in each trio in the case and control groups are shown in Fig 4B. Mean germline mutation rate estimated from observed autosomal MD in offspring of the case and control groups are shown in Fig 4C.

**Fig 4.**
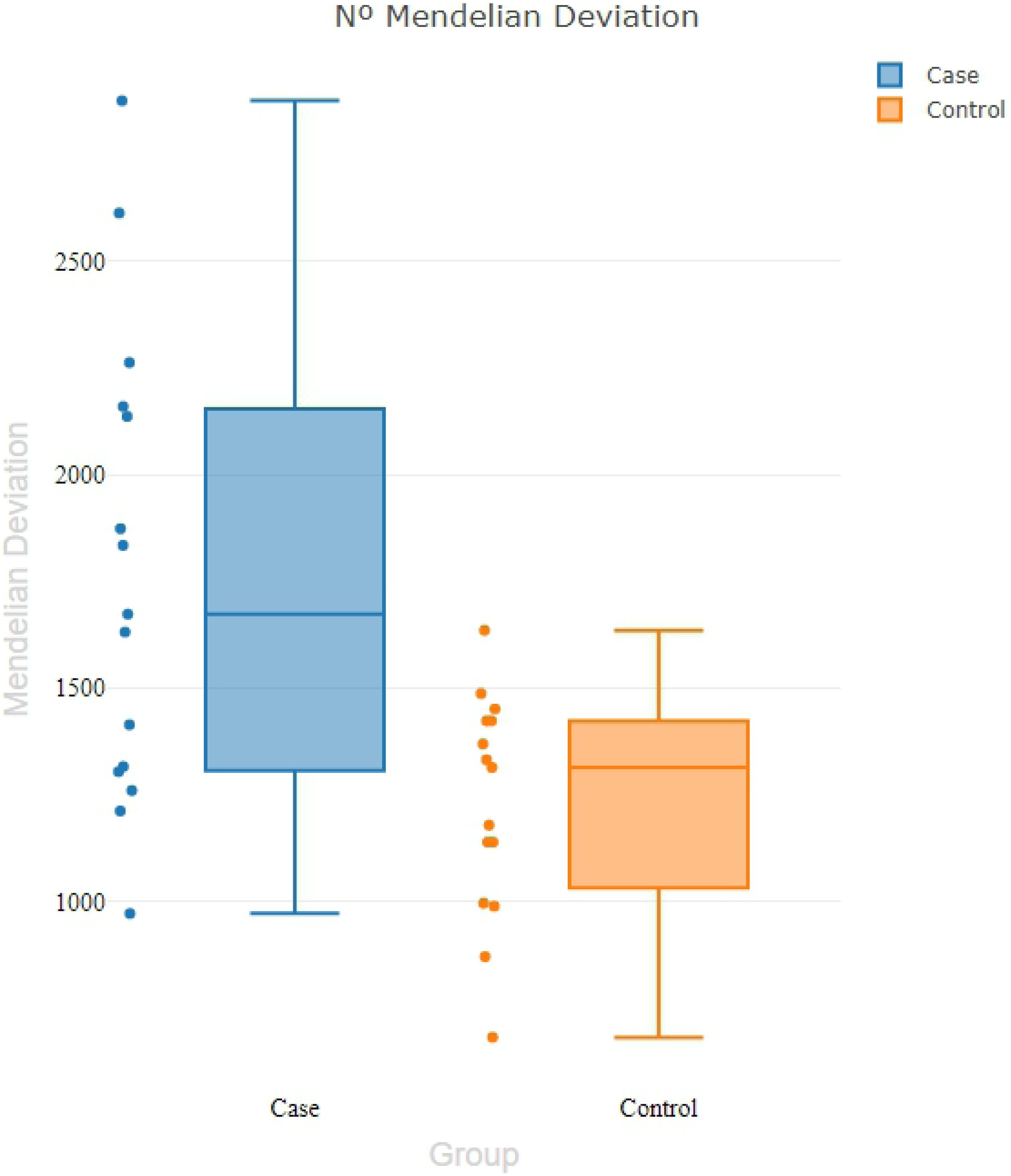
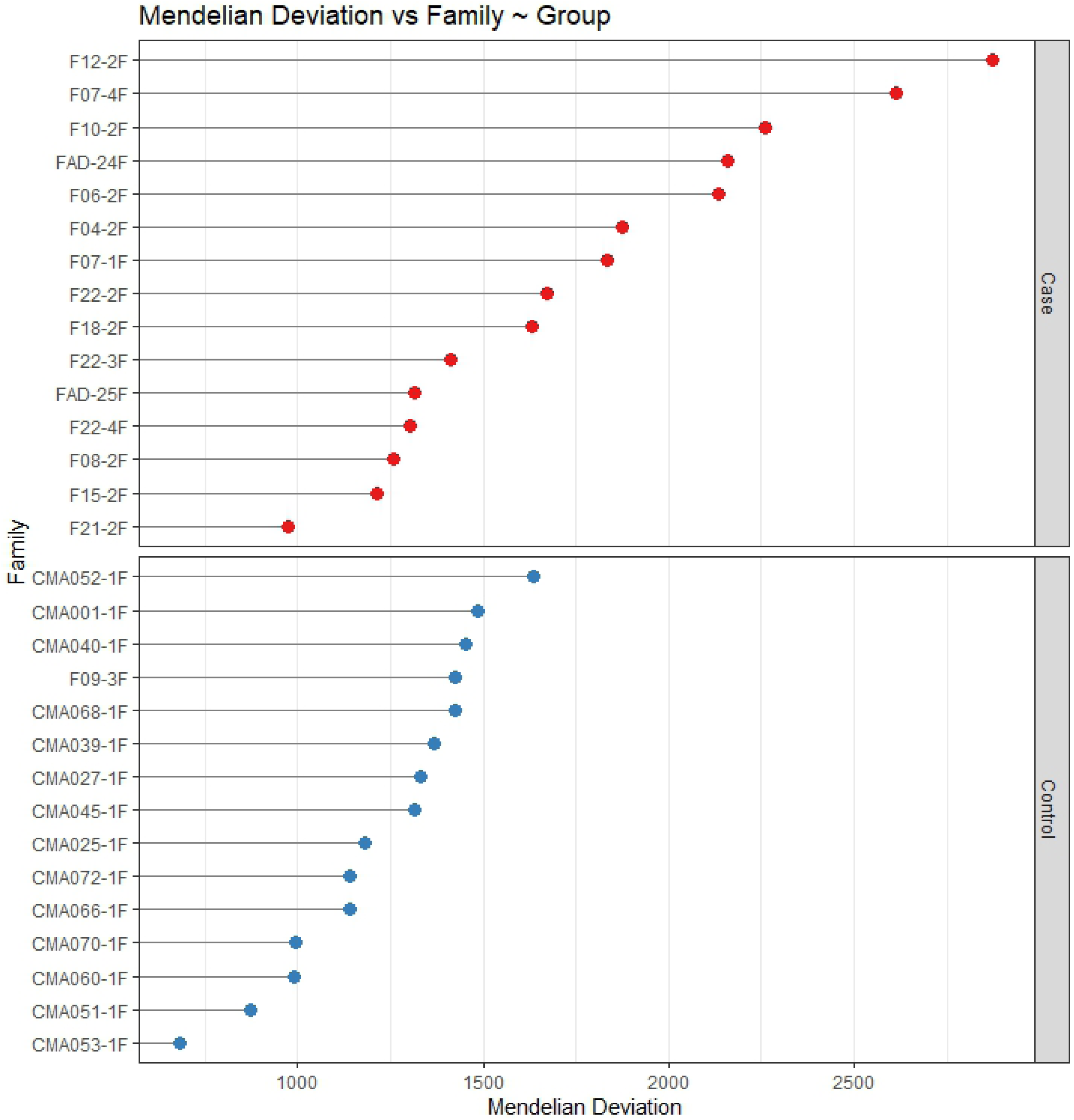
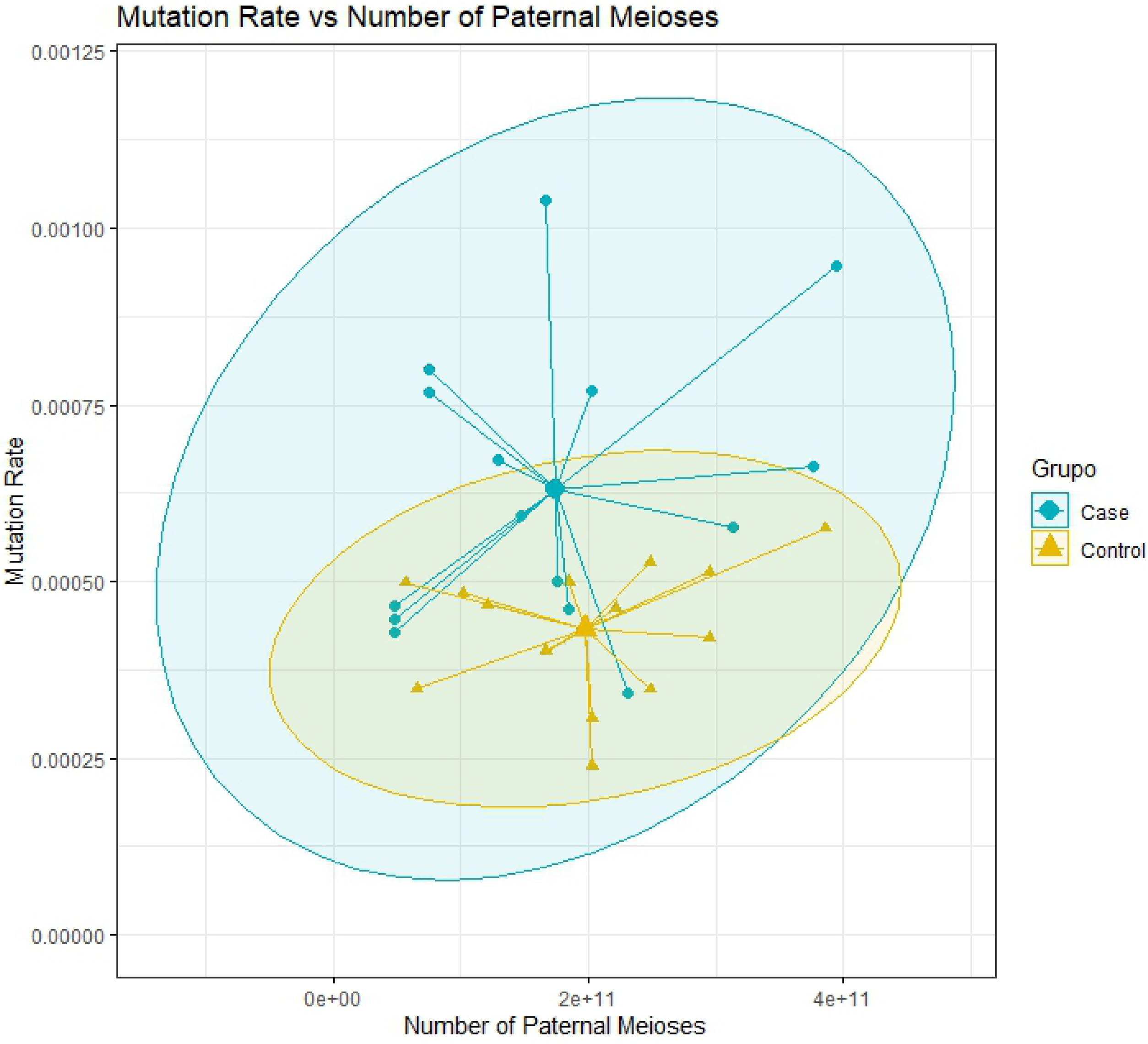
Representation of the variability of Mendelian deviations. **(A)** Mean numbers of Mendelian deviations in the offspring of the case and control groups. (B) Representation of the number of Mendelian deviations for each trio in the case and control groups. c) Mean germline mutation rates by MD in the autosomes of the progeny of the case and control groups.

The linear regression test (Fig 5) shows the result using the explanatory (Nρ!) and response (TM_DM_) variables. This model showed that the existing relationships between variables were not significantly explained: *p = 0*.31 and r^2^ = 0.12 (Fig 5A), and *p = 0.*38 and r^2^ = 0.06 (Fig 5B) for cases and controls, respectively. In this context, the difference between equations outlined by the regression model was used and resulted in y =13×10^−5^ + 2.6 (age). Thus, it is possible to conclude that MDs observed in the case group resulted from higher germline mutation rates than in the control group. Also, based on the age of paternal meiosis, MDs occurred 2.6 times as much in exposed than in non-exposed individuals.

**Fig 5.**
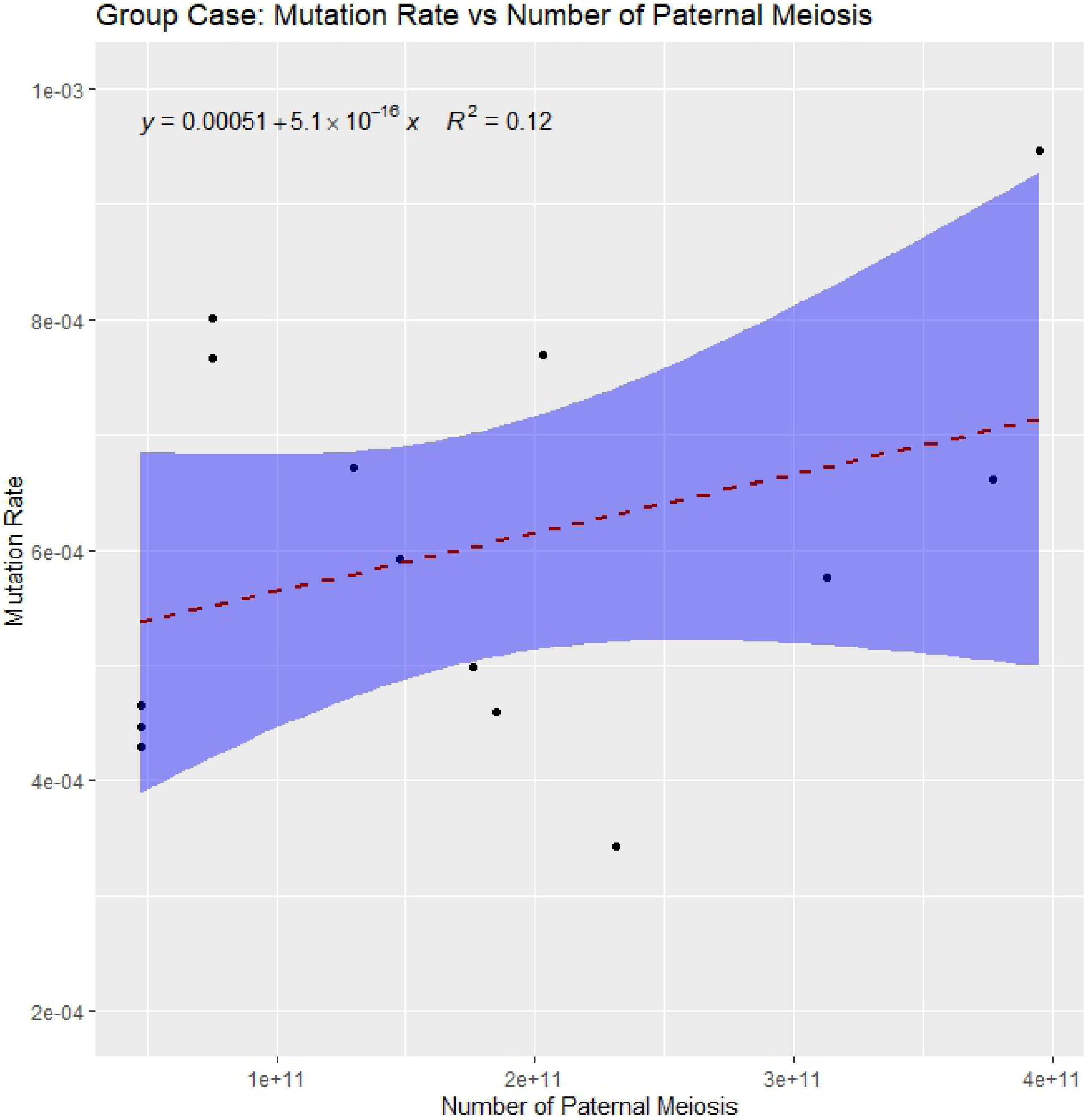
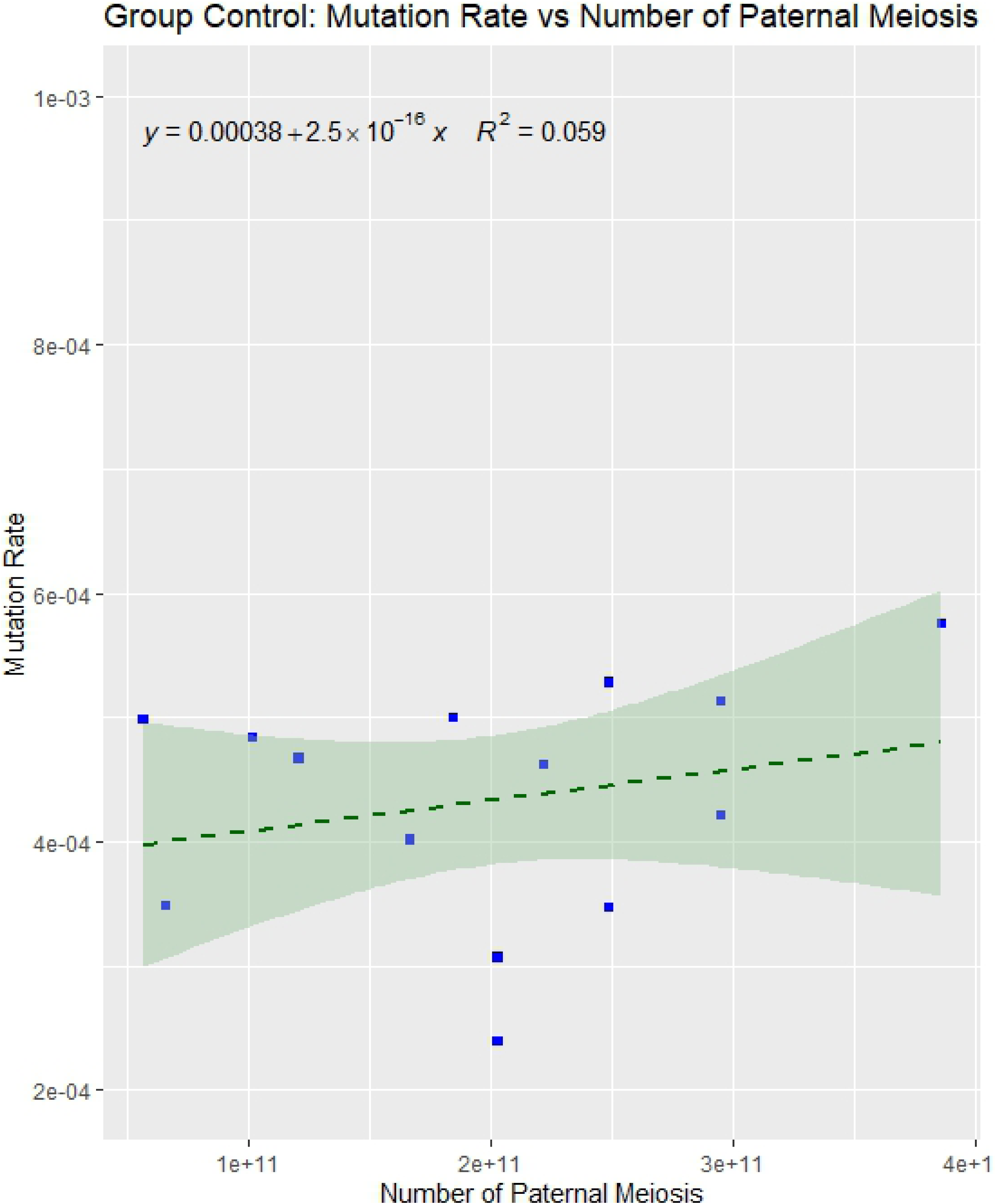
Linear regression models of TM_DM_ for the case (A) and control (B) groups.

Based on the linear regression observed *p-value*, and without considering the history of parental exposure, the small sample size could lead to an interpretation incurring type II errors. Cohen’s d was used to ameliorate this issue by evaluating the difference between the adjusted means, to measure the effect of sample size in this study. The estimated d-values were −2.15 and −3.09 for the case and control groups, respectively, and both were considered significant. In this context, the TM_DM_ was a biomarker sensitive enough to separate the group of children conceived and born to IR-exposed parents from controls conceived and born to non-exposed parents, given that comparisons were carried out in people from the same population with similar genetic background and admixture.

Analysis of variance was used to test the existence of equality in the mean number of paternal meiosis between the groups. Tukey’s multiple test (Fig 6) was used to estimate difference between means (Fig 6. A). Kernel density representation analyzing the probabilities of the findings in the samples, highlighting the difference between means is shown in Fig 6B. There was no significant difference between the mean number of paternal meiosis in the two analyzed groups (*t* = 1.238; p < 0.2).

**Fig 6.**
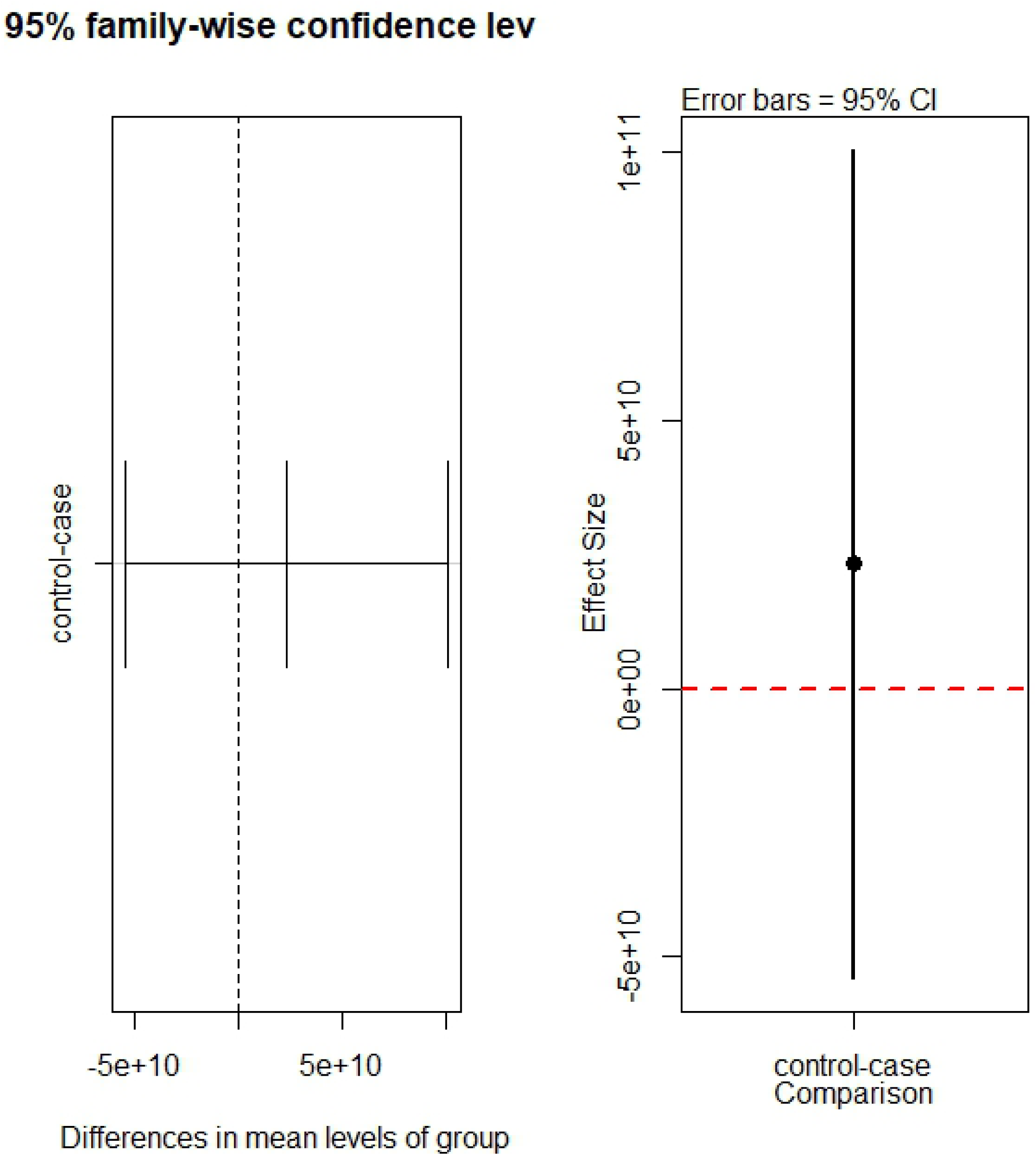
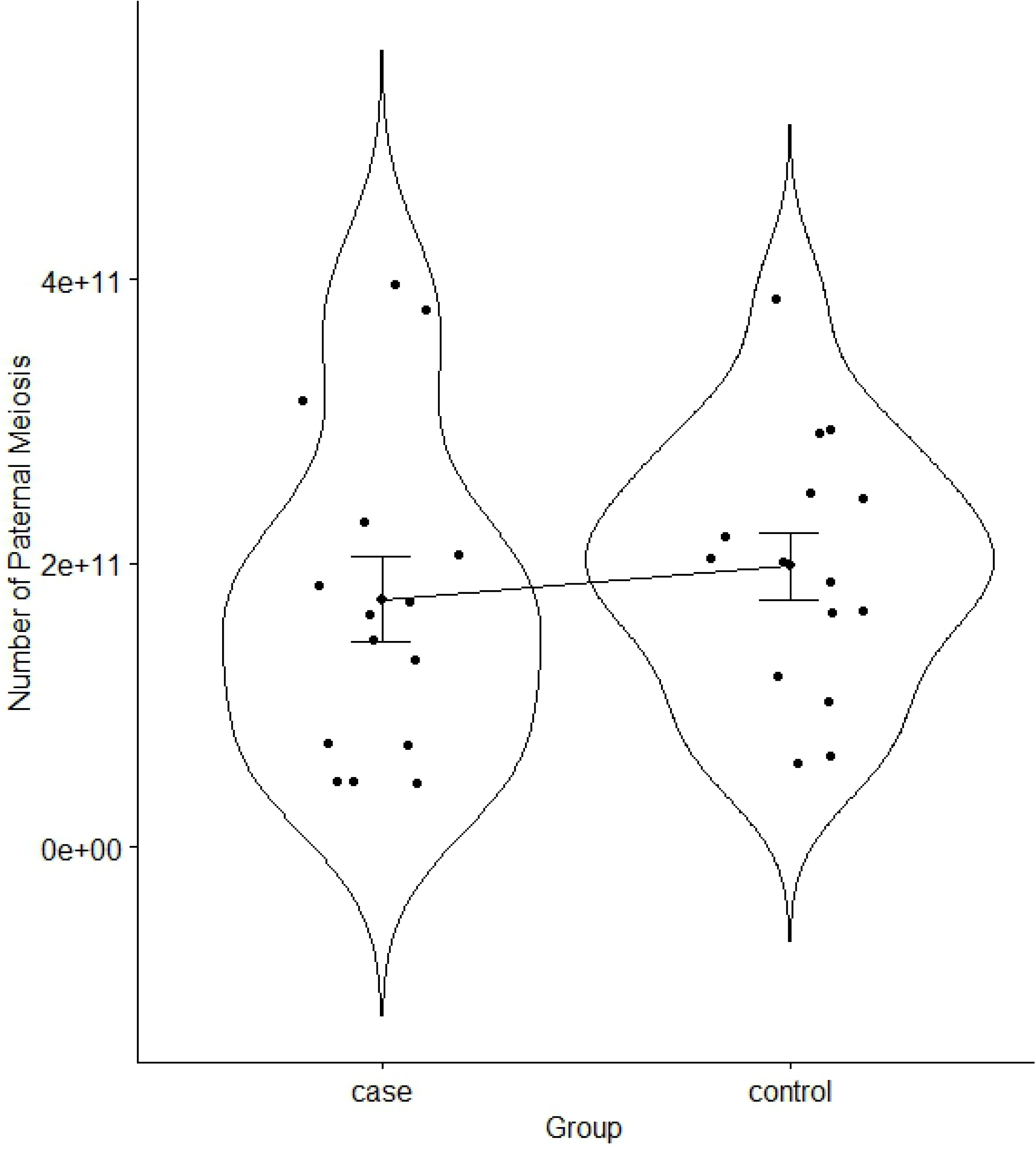
Analysis of average ages between groups. (A) The graphs represent theTukey’ s test. In this case, there was no difference between case and control means. (B) Kernel density representation analyzing the probabilities of the findings in the samples, highlighting the difference between means.

Some previously published studies on the types of DNA spontaneous base substitutions indicated all possible substitutions are well represented in germline cells [41]. Previous studies suggested that transition rates tend to be higher [42] than transversion rates [43]. The findings in the current study support these observations, since a higher proportion of transitions (74.5%) for cases compared to controls (25.5%) was observed in the children conceived after parental exposure to IR.

Procedures incorporated in SIPO identified 36% (9,522) and 26% (4,821) MDs whose origins could not be identified in the case and control groups, respectively, using previously established methods. In this context, SIPO can be used as an additional tool to define the parental origin of polymorphic variants obtained from SNP array genotypes.

To validate the findings of the current study, which analyzed the TM_DM_ of a small cohort of children conceived after their parents were accidentally exposed to ionizing radiation from Cs-137, the authors recommend the development of studies involving larger cohorts. It might be advisable to include a wider range of absorbed doses, resulting from either therapeutic or occupational exposure, to assess the real potential of using observed Mendelian deviations as retrospective biomarkers for IR exposure in human populations. In the present study, as the case and control groups belonged to the same population, and therefore were subjected to similar general environmental effects, the significant TM_DM_ differences observed between the case and control groups were most likely due to parental exposure to IR. Thus, it is safe to assume that low doses of low-LET radiation can induce MDs that can be identified and quantified and therefore used as a biomarker to study human populations according to their history of exposure to mutagenic insults.

## Conclusions

This study pioneered the analysis of MDs observed in parent-offspring trios as biomarkers of exposure to low doses of ionizing radiation. We succeeded in retrospectively estimating the rate of germline mutation in a human population accidentally exposed to low doses of radiation from Cs-137 and estimated the burden of germline mutations in the offspring. In summary, there was a 46.5 % increase in the TM_DM_ of the F1 generation of parents accidentally exposed to Cs-137 IR, from a radiological accident in Goiânia, with absorbed doses ranging from 0.2 to 0.5 Gy. Therefore, we conclude that TM_DM_ is a potentially useful biomarker to estimate parental exposure to IR and for human biomonitoring. In this context, future studies involving the behavior of MDs in genomic and mutagenic hazards, caused by exposure to environmental agents, may provide important knowledge of the biological effects, risks and mechanisms resulting from human exposure to such agents.

Finally, we are confident SNP array data can be used to estimate ionizing radiation-induced mutagenesis in human populations, provided the appropriate bioinformatics and statistical tools are used to extract the necessary information for biological inferences and to validate the scientific hypotheses underlying each investigation.

## Acknowledgements

The authors wish to thank the voluntarily participation of the patients. This study was supported by CNPq (Conselho Nacional de Desenvolvimento Científico e Tecnológico), FAPEG (Fundação de Amparo à Pesquisa do Estado de Goiás), CAPES (Coordenação de Aperfeiçoamento de Pessoal de Nível Superior), D.M.S. and A.D.C. wish to thank for their scholarships. We also wish to express gratitute to CARA (Centro de Assistência ao Radioacidentado da SES-GO) for helping with contacting the group of exposed parents. Moreover, we thank Dr. Fernando Nodari for assisting with the issues from the software ChAS^®^ (Affymetrix, USA). Lastly, we thank the volunteers who selfishly agreed to participate in the study. The funders had no role in study design, data collection and analysis, decision to publish, or preparation of the manuscript. A.R.C. is a CNPq research fellow.

